# DETERMINATION OF CYTOTOXICITY AND FUNCTIONAL GROUPS IN THE RIGHT-WING OF MUSCA DOMESTICA

**DOI:** 10.1101/2025.11.11.687935

**Authors:** Rommel Osabel, Princess C. Domingo, Jea Farida R. Guroalim, Leonel P. Lumogdang

**Affiliations:** Teacher Education Program, Institute of Teacher Education and Information Technology, Southern Philippines Agri-business and Marine and Aquatic School of Technology (SPAMAST), Malita Davao Occidental. 8012 Philippines; Research and Laboratory Services Center, Southern Philippines Agri-business and Marine and Aquatic School of Technology (SPAMAST), Malita Davao Occidental, 8012 Philippines

**Keywords:** *Musca domestica*, Bioactive compounds, Cytotoxicity, BSLA

## Abstract

This study focused on determining the cytotoxicity of right-winged *Musca domestica* and assessing its bioactive compounds using FTIR spectroscopy. The cytotoxicity was evaluated using the Lc50 of the brine shrimp lethality assay. The Brine Shrimp Lethality Assay (BSLA) result shows that the extract obtained using the aqueous extract has the highest cytotoxicity. The FTIR analysis results showed different characteristic peaks and confirmed the presence of a functional group in the extract obtained using the aqueous extracts of Right-wing *Musca domestica*.

## INTRODUCTION

Houseflies are considered disease vectors due to their hopping and feeding behavior in their pathogenic substrates (*Nazari et al., 2017*). This two-winged insect is known to survive in pathogenic communities, which exposes them to a wide range of pathogens. This has led them to develop effective innate immune systems that rapidly respond to environmental threats, such as pathogens (Szpila & Pape, 2008). The wings of insects, although often overlooked in chemical investigations, can be reservoirs of secondary metabolites that contribute to their survival strategies. The insect wings’ resistance to microbial infestation, developed over 400 million years of evolution, is evidenced by surface nanopillar topography, which dually repels fungal conidia and kills bacterial cells upon direct contact (*Ivanova et al., 2021*)

Numerous studies have been carried out to identify which wing possessed the ailment and the cure. The right wing harbors the cure, and the left wing possesses the ailment. The right wing has a microbial community that efficiently inhibits pathogens from their substrates (Atta, 2014; *Claresta et al., 2020; Asril et al., 2021*). However, the aforementioned studies did not determine the cytotoxicity and functional groups in the right wing responsible for inhibiting pathogens in their substrates.

Hence, this study focused on the determination of cytotoxicity and functional groups in the right wing of *Musca domestica* using Brine Shrimp Lethality Assay (BSLA) for cytotoxic profiling and Fourier-transform infrared spectroscopy (FTIR) for functional group analysis. BSLA is a well-established method for preliminary bioactivity assessment, and it is successful in cytotoxicity screening (*Balinado et al., 2019*). FTIR has been proven effective in confirming bioactive functional groups (Karpagasundari et al., 2014). By employing the aforementioned methods in the right wing of houseflies, the researchers aim to provide data on the cytotoxicity and functional groups in the right wing of *M. domestica*, shedding light on their potential ecological functions and the broader significance.

## MATERIALS AND METHODS

### Research Locale

The fly samples were collected randomly. This study followed sustainable fly collection procedures. Prior to the commencement of the data-gathering procedure in Santa Maria, Davao Occidental, Philippines. All the necessary permits and letters were obtained from school authorities and environmental management departments. These permissions include authorization from the Department of Natural Resources (DENR) and authorization to use the school laboratory and the hatching facility. In addition, ethical guidelines on insect sampling were followed.

### Collection and Cleaning of Samples

The researchers used an improvised trap to capture flies. The main body of the fly trap was made out of a fine net cylinder, about 2 feet tall, and a diameter of about 1.2-1.5 feet. Baits, preferably carrion/animal remains, were utilized to lure house flies. Within each selected area for sample collection, only the desired species of fly that happened to fall into the traps were utilized for the study. We based our selection on an article by Medhekar (2015) to identify the desired fly species. The researchers performed a dichotomous key for species identification. As a reference for the traits of stable flies, the researchers utilized the study by *Rochon et al. (2021)*. Traps made out of water bottles were also utilized.

**Figure 1.**
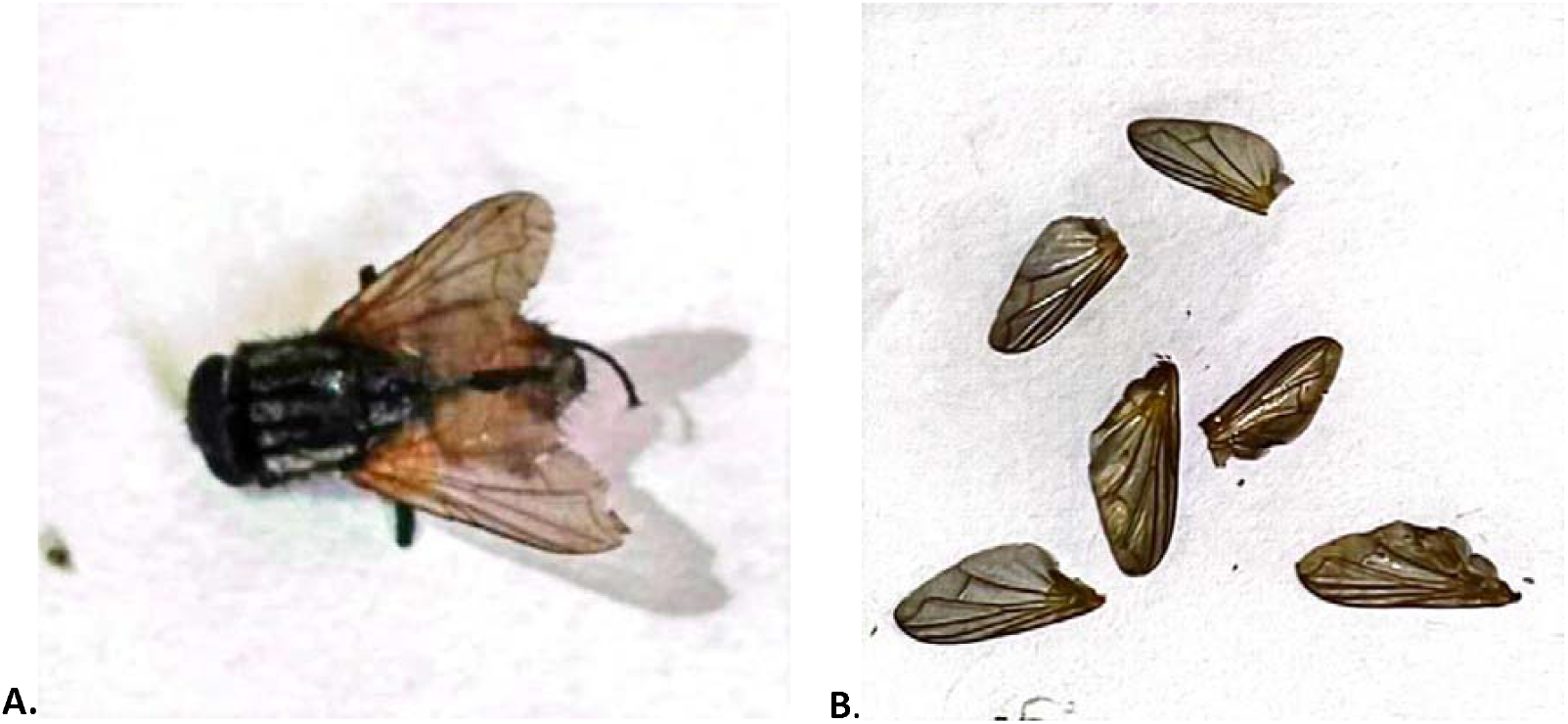
shows the whole image of Musca domestica(A) and the Right Wing(B) of the Musca domestica used in the study

The weight of the collected housefly ranges from 0.005 to 0.007 grams, while the body size ranges from 6-7 millimeters. The right wing of the housefly used in the study ranges from 5-7 millimeters; however, the actual weight of the individual wings was not determined in the study due to the technical limitations of the laboratory facilities. Moreover, the age of the housefly was not determined in the study due to the lack of an instrument in the facilities that could determine the age of the housefly.

In total, 2.25 grams of wing samples were collected, yielding 724,500 μg (0.7245 grams) of crude extract. The extraction of compounds and BSLA lasted 1 week and 3 days. The first week was used for the extraction of compounds. The first day of the second week was used to hatch the brine shrimp cysts. The brine shrimp containers and saline solution were prepared, the extracts were serially diluted, and treatments were applied on the second day of the second week.

The fly samples lured by the traps were kept in a container to keep them alive. The wing samples were processed immediately. A dissecting kit separated the wing samples from the insect’s thorax. Amputation was performed with the aid of a dissecting microscope. The wing samples were powdered and processed in the SPAMAST Research and Laboratory Services Center.

### Preparation of extracts

The compounds in the right wings of *Musca domestica* were extracted using ethanolic, aqueous, and 50:50 extraction methods. A varied mass of fly wings was mixed into a fixed amount of solvent for each extraction method. The ethanolic extract was prepared by grinding wing samples into powder using a mortar and pestle. The 750 mg powdered wing samples were used. The researchers measured 50 mL of absolute ethanol and poured it into a 100 mL Erlenmeyer flask. The researchers then put 750 mg of the powdered wing sample in the Erlenmeyer flask containing the extraction solvent. The resulting mixture was agitated to facilitate the extraction. The flask was sealed with a cork. The wing samples were soaked in absolute ethanol for 24 hours. The mixture was labeled with relevant information and stored in a reagent safe. The solvent was then evaporated using a water bath until only the crude extract was left in the Erlenmeyer flask.

The researchers pounded the wing samples into powder using a mortar and pestle to prepare the aqueous extract. 750 mg powdered wing samples were used. The researchers measured 50 ml of distilled water and poured it into a 100 ml Erlenmeyer flask. The researchers then put 750 mg of the powdered wing sample in the Erlenmeyer flask containing the extraction solvent. The resulting mixture was agitated to facilitate the extraction. The flask was sealed with a cork. The wing sample was soaked in distilled water for 24 hours. The mixture was labeled with relevant information and stored in a reagent safe. After 24, the mixture was filtered to separate the liquid and solid portion of the mixture. The solvent was then evaporated using a water bath until only the crude extract was left in the Erlenmeyer flask.

To prepare the blending of the ethanolic and aqueous extract, the researchers grind the wing samples into powder using a mortar and pestle. 750 mg powdered wing samples were used. The researchers measured 25 mL of distilled water and 25 mL of absolute ethanol. The researchers then poured the solution into a 100 mL Erlenmeyer flask. The researchers then put 750 mg of the powdered wing sample in the Erlenmeyer flask containing the extraction solvent. The resulting mixture was agitated to facilitate the extraction. The flask was sealed with a cork. The wing sample was soaked in distilled water for 24 hours. The mixture was labeled with relevant information and stored in a reagent safe. After 24, the mixture was filtered to separate the liquid and solid portion of the mixture. The solvent was then evaporated using a water bath until only the crude extract was left in the Erlenmeyer flask. Each of the extracts yielded at least 241.50 mg of crude extract. Combining all 3 extracts from each extraction method yielded 724.50 mg crude extract. The researchers prepared four solutions with varied concentrations (1000 ppm, 500 ppm, 100 ppm, and 10 ppm).

### Brine Shrimp Lethality Assay (BSLA) analysis

The brine shrimp cysts (eggs) were purchased from the local supplier. The researchers created a saline solution in a brine shrimp hatchery. In a 2 L plastic bottle, 1 gram of *Artemia salina* cyst was soaked in 500 mL of natural water for 1 hour. 17 grams of sea salt were dissolved in the 500 mL natural fresh water. The recommended salinity for hatching brine shrimp cysts is around 25-35 parts per thousand (ppt). The saltwater solution was aerated with an air pump to provide oxygen and hold the cyst in a suspension. The researchers waited 24 hours for the cysts to hatch. During that period, the cyst developed into the first larval stage of its development (nauplii).

The researchers collected the nauplii using a micropipette and transferred the collected larva to a beaker with saline solution. The researchers prepared 45 sauce cups to be used as the brine shrimp containers. They filled the cups with saline solution and added 10 brine shrimp to each cup, adding three drops of dimethyl sulfoxide (DMSO) (*Lumogdang et al., 2021*).

### Preparation of treatments

The 45 cups were divided into 3 groups of 15. Group 1 was labeled T1 (Solvent 1), group 2 is T2 (Solvent 2), and group 3 is T3 (Solvent 3). Group 1 was treated with the crude extract obtained by the aqueous extraction method. Group 2 was treated with the extract obtained by the 50:50 extraction method. Group 3 was treated with the extract obtained by the ethanolic extraction. For each group, 12 cups received the experimental treatment. The 12 cups were divided into 4 groups of 3. Each group will be treated with the exposure solutions with varying concentrations (1000 ppm, 500 ppm, 100 ppm, and 10 ppm) of their respective extracts. The remaining 3 cups out of 15 were the control group. Apply the treatments and wait for 24 hours.

The researchers inspected the cups after 6, 12, and 24 hours using a magnifying glass and counted the number of nauplii that survived. By calculating the mortality rate, the researchers determined the LC_50_ concentration using probit analysis.

### FTIR Analysis

The liquid extracts of the right-wing *Musca domestica* were sent to the Analytical Science laboratory of the University of Santo Tomas, Manila, Philippines.

### Data Analysis

The brine shrimp lethality assay (BSLA) result was interpreted based on the papers of *Guia et al*. (2025, Lumogdang et al. (2021), and *Meyer et al*. (1982). FTIR spectral data were interpreted based on the protocol of *Kumar et al*., 2019.

## RESULTS AND DISCUSSIONS

Table 1 shows the cytotoxicity of the right-wing of *Musca domestica* extracts obtained using aqueous, Mixed extract of Aqueous and Ethanolic, and ethanolic extracts after 6, 12, and 24 hours of exposure to Brine shrimps. Brine shrimps were tested in flies’ right-wing extract obtained using three different extracts with four concentrations—10 ppm, 100 ppm, 500 ppm, and 1000 ppm. The toxicity values (LC50) were computed and described using the Clarkson Toxicity Index used by *Guia et al*. (2025).

**Table 1.**
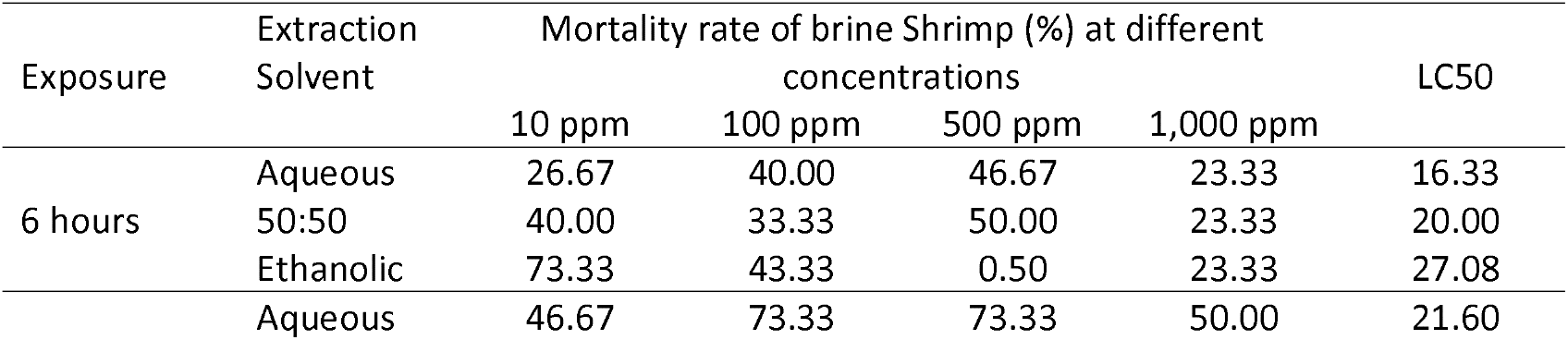
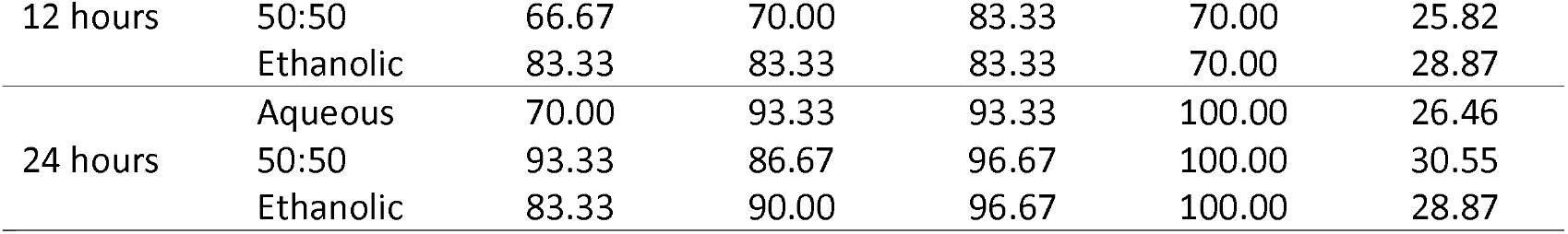
Cytotoxicity of right-wing *Musca domestica* after 6 hours, 12 hours, and 24 hours.

After 6 hours, the brine shrimp showed reactions to the concentrations. The wing extract obtained from the aqueous extraction method, 10 ppm, resulted in a toxicity value of 16.33. The extract from the 50:50 extraction method, 10 ppm, induced a toxicity value of 20.00 and is interpreted as toxic. Moreover, the ethanolic extract has a toxicity value of 27.08 ppm and is interpreted as highly toxic. After 12 hours, the cytotoxicity of wing extracts was obtained using aqueous, 50:50, and ethanolic wing extraction methods. The aqueous extract has a toxicity value of 21.60 ppm, which is interpreted as toxic. The 50:50 extract of Aqueous and ethanolic obtained using the 50:50 extraction method induced toxicity of 25.82 ppm and is interpreted as toxic. The ethanolic extract has a toxicity value of 28.87 ppm and is described as toxic. After 24 hours, the aqueous extract had a toxicity value of 26.46 ppm and was interpreted as toxic. The mixed Aqueous and Ethanolic extract had a toxicity value of 30.55 and was interpreted as toxic. The ethanolic extract has a toxicity value of 28.87 and is interpreted as toxic.

The aqueous extraction method for the right wing of *M. domestica* yields the highest toxicity value compared to other extraction methods, indicating its potential for extracting bioactive compounds. The brine shrimp lethality assay effectively and conveniently detects bioactive compounds, as it strongly correlates with cytotoxicity and other biological activities (*Meyer et al*., 1982). The aqueous extract with the highest toxicity value was subjected to FTIR analysis.

Table 2 shows the FTIR-ATR spectra results for the aqueous solution (R1), revealing a prominent peak at 1639.49 cm-1 and 3329.14 cm^-1^, indicating the presence of distinct molecular phenomena. The peak value, 1639.49 cm^-1^, corresponds to the stretching vibrations of C=O bonds and saturated and conjugated C=C bonds. These molecular vibrations indicate the presence of amides and alkenes functional groups within the sample, respectively. The peak value of 3329.14 cm-1 corresponds to the stretching vibrations of O-H bonds and (2) two N-H stretches. These molecular vibrations indicate the presence of alcohol, amines, and amide functional groups within the sample.

**Table 2.** FTIR spectra of the aqueous extract of right-wing *Musca domestica*.

| FUNCTIONAL GROUPS |  |  |  |
| --- | --- | --- | --- |
| REPLICATES | PEAK VALUE<br>(cm<sup>-1</sup>) | PHENOMENO<br>N | FUNCTIONA<br>L GROUP |
| Aqueous Extract<br>(R1) | 1639.49 | C=O stretch | Amides |
|  |  | C=C stretch (sat) | Alkenes |
|  |  | C=C stretch<br>(conj) | Alkenes |
|  | 3329.14 | O-H stretch | Alcohols |
|  |  | N-H stretch | Amines |
|  |  | N-H stretch | Amides |
| Aqueous Extract<br>(R2) | 1631.78 | C=O stretch | Amides |
|  |  | C=C stretch (sat) | Alkenes |
|  |  | C=C stretch<br>(conj.) | Alkenes |
|  | 3350.35 | O-H stretch | Alcohols |
|  |  | N-H stretch | Amines |
|  |  | N-H stretch | Amides |
| Aqueous Extract<br>(R3) | 1631.78 | C=O stretch | Amides |
|  |  | C=C stretch (sat) | Alkenes |
|  |  | C=C stretch<br>(conj) | Alkenes |
|  | 3354.21 | O-H stretch | Alcohols |

In aqueous solution (R2), revealed a prominent peak at 1631.78 cm^-1^ and 3350.35 cm^-1^, indicating the presence of distinct molecular phenomena. The peak value 1631.78 cm^-1^ corresponds to the stretching vibrations of C=O bonds and saturated and conjugated C=C bonds. These molecular vibrations indicate the presence of amides and alkenes functional groups within the sample, respectively. The peak value of 3329.14 cm-1 corresponds to the stretching vibrations of O-H bonds and (2) two N-H stretches. These molecular vibrations indicate the presence of alcohol, amines, and amide functional groups within the sample.

Lastly, an aqueous solution (R3) revealed a prominent peak at 1631.78 cm^-1^ and 3350.35 cm^-1^, indicating the presence of distinct molecular phenomena. The peak value 1631.78 cm^-1^ corresponds to the stretching vibrations of C=O bonds and saturated and conjugated C=C bonds. These molecular vibrations indicate the presence of amides and alkenes functional groups within the sample, respectively. The peak value of 3354.31 cm-1 corresponds to the stretching vibrations of O-H bonds. These molecular vibrations indicate the presence of an alcohol functional group within the sample.

The current study result supports previous studies conducted by *Claresta et al.* (2020), which found that the right wing of *Musca domestica* can inhibit the growth of Escherichia coli in contaminated drinks. Moreover, it also supports the result of Atta (2020), which found that all the media cultivated with right-wing extract were free of bacterial and fungal growth; however, microbial growth was observed on the left wing of Musca domestica. The high cytotoxicity of the three extracts could be attributed to the bioactive compounds present in the right wing of *Musca domestica*, namely Antibacterial peptide MD7095(*Lu et al.*, 2006; Asril et al.,2021), Antimicrobial peptide Cecropin (Mdc) (*Lu et al*.,*2012: *Asril et al.*, 2021*), Antimicrobial peptide (MDAP-2) (*Pei et al.*, 2014; *Asril et al.*, 2021), Antifungal peptide (MAF-1) (*Fu et al., 2009; Asril et al., 2021*), 1-lysophosphatidylethanolamine (*Meylaers et al., 2004; Asril et al., 2021*), Phenylacetaldehyde (*Ali et al., 2018; *Arora et al.*, 2011: Asril et al., 2021*), Bacteriophage and Adenosine monophosphate(*Niode et al; 2022*). The FTIR detection of alkene and amines in the ethanolic extract of the right-wing of *Musca domestica* supports the functional groups present in the protein nature of the following bioactive compounds on the Antibacterial peptide MD7095, Antimicrobial peptide Cecropin, Antimicrobial peptide (MDAP-2), Antifungal peptide (MAF-1). The bioactive compounds in the right wing of Musca domestica could be attributed to the bacterial communities present in its wings. Previous studies isolated and identified the bacterial communities of the right-wing of Musca domestica found the following bacterial species: *Micrococcus luteus, Sphingobacterium sp., Bacillus subtilis, Staphylococcus aureus, S. xylosus, Acinetobacter spp., Brucella melitensis, Escherichia coli, Klebsiella oxytoca, Proteus vulgaris, P. fuorescens* that are responsible for the production of abundant bioactive compounds (*Laziz et al.*, 2021; *Kanan et al.*, 2020). The presence of these bacterial communities contributes to the inhibition of pathogens from their substrate.

Moreover, the FTIR detected the Alcohol and Amine functional group in the bioactive compound 1-lysophosphatidylethanolamine. Moreover, the FTIR also detected alkene, which supports the presence of the R-group in the bacteriophage. However, the carbonyl group in phenylacetaldehyde was not detected in the aqueous extract, which may be due to the inability of the solvent to extract the carbonyl-based bioactive compounds. It may be possible to be detected by the other solvent, but only the aqueous extract of the right wing of the *Musca domestica* was subjected to FTIR. The aqueous extract did not detect the ester-based bioactive compounds in Adenosine monophosphate (AMP).

The study’s findings strongly support the medicinal properties of the right-wing *Musca domestica*. However, comparative studies must be done to evaluate the presence of bioactive compounds in different maturity stages of *Musca domestica* and determine the maturity of Musca domestica that can be used for medicinal purposes.

## CONCLUSION

The Brine Shrimp Lethality Assay (BSLA) determined that all extracts from the three extraction methods were toxic, with aqueous extracts showing the highest toxicity. This cytotoxicity of the right-wing may correlate with the previously observed biological activities of the right-wing and larvae of *M. domestica*. Moreover, active functional groups, namely amides, alcohol, amines, and alkenes, confirmed the previously isolated bioactive compound in the right wing of *Musca domestica*. The presence of active bioactive compounds in the right wing of Musca domestica is a promising result as a potential bioactive source for drug discovery and development. The study’s findings favor the utilization of *Musca domestica* as a source of bioactive, which is also an environmentally friendly and preventive measure to control *Musca domestica* in the environment.

## ACKNOWLEDGEMENT

The researchers wish to acknowledge the SPAMAST Research and Laboratory Services Center (RLSC) for allowing them to use its laboratory facilities to implement the study.

